# Inferring spatial gene expression from tissue images using large-scale histology foundation model with SpaFoundation

**DOI:** 10.1101/2025.08.07.669202

**Authors:** Ning Zhang, Yahui Long, Sibo Xia, Pengzhen Jia, Yujie Jin, Siqi Chen, Jinmiao Chen, Min Li

## Abstract

Spatial transcriptomics (ST) has revolutionized biological research by enabling the joint profiling of gene expression and spatial context, along with histological images. However, current existing ST technologies remain high-cost and time-consuming, hindering their broader clinical applications. Although computational methods have been developed to infer gene expression directly from histology images, these methods still suffer from limited accuracy and spatial resolution due to insufficient training data and model capacity. Here, we introduce SpaFoundation, a large-scale histology foundation model designed to accurately predict spatial gene expression from tissue images. SpaFoundation employs a teacher-student Vision Transformer (ViT) architecture to learn generalizable histological representations by modeling potential dependencies among image patches. Notably, we incorporated self-distillation and masked image modeling (MIM) jointly to capture high-level semantic representations and fine-grained structural features, enriching spots’ representations. The model is pretrained on 1.79 million patches spanning 26 tissue types, with 80 million parameters. We validated SpaFoundation using 117 samples, demonstrating its flexibility across different spatial resolutions and superior performance in spatial gene expression prediction, as well as strong transferability to downstream tasks such as tumor detection and spatial domain clustering. Our results highlight the potential of large-scale foundation model to learn informative histological representations and underscore the benefits of domain-specific pretraining in extracting task-relevant representations, paving the way for foundation model-driven spatial gene expression inference. The implementation and pre-trained weights of SpaFoundation are publicly available at https://github.com/NingZhangCSUBio/SpaFoundation.

## 1 Introduction

Spatial transcriptomics (ST) has emerged as a transformative technology that simultaneously captures gene expression and spatial localization information within intact tissues^1–4^, enabling the exploration of spatial gene expression patterns, cell-cell communications, and the spatiotemporal dynamics of tissue development^5–8^. Existing ST technologies can be roughly divided into two categories, including sequencing-based approaches (e.g., STARmap^9^, 10x Visium, SLIDE-seqV2^10^, and Stereo-seq^11^) that offer genome-wide coverage at relatively coarse spatial resolution, and imaging-based approaches (e.g., 10x Xenium, DNA seqFISH+^12^, MERSCOPE^13^, and Nanostring CosMx^14^) that provide high spatial resolution at limited gene throughput. However, both of them remain high-cost and time-consuming, hindering broader applications.

Previous studies have demonstrated that histology image and gene expression profiles are intrinsically linked and both reflect the underlying biological state and spatial organization of tissues, suggesting the possibility to predict spatial gene expression from histology images^15,16^. Motivated by this observation, computational approaches have been developed to infer spatial gene expression profiles from histology images, such as His2ST^17^, His2Gene^18^, THItoGene^19^, BLEEP^20^, TRIPLEX^21^, and GHIST^22^. These existing methods typically employ either convolutional neural network (CNN) or Transformer architectures to extract visual features around captured spots from histology image patches, which are then utilized to capture biologically relevant gene patterns. For example, ST-Net utilized a convolutional neural network to extract histological features from images, and then applied a fully connected layer to predict gene expression of spots^15^. His2ST combined CNN and transformer to extract multi-view spatial features from histology images for improved prediction^17^. His2Gene adopted a vision transformer to learn histological representations for the prediction^18^. Some methods combine histology image with gene expression during training to boost performance. His2Gene, for example, learned a shared embedding space aligning image and gene expression modalities, whereas BLEEP employed contrastive learning to construct joint image-expression embedding for more robust prediction^20^. Despite their promising results, these methods often suffer from limited generalizability across tissues, organs, or sequencing platforms, as their architectures are typically tailored to specific datasets and tasks. Moreover, constrained model capacity and insufficient training data, typically confined to single tissue slice, restrict their ability to capture intricate relationships between tissue morphology and gene expression patterns.

Recently, an emerging trend has focused on leveraging pathology foundation models to extract histological features for improved gene expression prediction. For instance, FmH2ST utilized a pretrained foundation model from computational pathology to learn morphological representations^23^. TRIPLEX adopted a ResNet18^24^ backbone to capture multi-resolution histological features that enhance prediction accuracy^21^. HISTEX employed UNI^25^ as its backbone histological feature extractor^26^. While these methods have made significant advancements, they still face inherent limitations. The foundation models they rely on are typically pretrained on pathology images, which differ fundamentally from hematoxylin-and-eosin (H&E)-stained images in spatial transcriptomics. Unlike pathology images that primarily capture cellular and structural morphology but lack direct molecular information, spatial transcriptomics H&E images are intrinsically linked to the underlying molecular landscape by measuring gene expression at defined spatial coordinates, possessing richer contextual features for accurate gene expression inference. However, large-scale histological foundation models specifically tailored and pretrained on spatial transcriptomics H&E images remain largely unexplored.

In this study, we introduce SpaFoundation, a histology foundation model to infer spatial gene expression. SpaFoundation constructs a teacher-student Vision Transformer architecture as basic framework, aiming to learn general-purpose histological representations from large-scale curated image datasets by modeling intrinsic dependencies between image patches. To efficiently capture both high-level semantic context and fine-grained structural details, we integrated self-distillation and masked image modeling (MIM) into the pretraining process. We pre-trained the model on HEST-1K-3 dataset, comprising 1.79 million image patches from 1,113 samples across 26 tissue organs. Evaluated on 117 samples spanning multiple technological platforms, SpaFoundation consistently outperformed 10 state-of-the-art traditional methods, achieving superior or comparable performance under minimal or zero-shot fine-tuning across four downstream tasks, including spatial clustering, tumor detection, spatial gene expression prediction, and high-resolution gene expression inference. Furthermore, ablation studies validated the model’s ability to extract biologically meaningful and transferable features from histology images. Collectively, these results position our SpaFoundation as a pioneering step toward a new paradigm of foundation model-driven spatial gene expression inference.

## 2 Methods

### 2.1 Datasets for SpaFoundation pretraining

We adopt HEST-1K (curated by Guillaume et al.) as pretraining dataset for SpaFoundation^27^. HEST-1K encompasses spatial transcriptomics data generated using three distinct technologies, i.e., Spatial Transcriptomics, Xenium, and Visium (including Visium HD), with each sample paired with a corresponding histological image. The dataset includes 1,229 samples from 153 public or internal cohorts, spanning two species: Homo sapiens (47.2%) and Mus musculus (52.8%). After tissue detection and segmentation, HEST-1K yields approximately 2.1 million expression–morphology pairs (each patch sized 224×224 pixels) and 76.4 million nuclei. Notably, three datasets used in the downstream tasks, i.e., HER2+^28^, cSCC^29^, and HBCIS^15^, are excluded from pre-training. The remaining subset, referred to as HEST-1K-3, includes 1,113 samples across 26 distinct tissue types.

### 2.2 Problem formulation

SpaFoundation aims to infer spatial gene expression of spots from given H&E-stained histopathology image *I* ∈ ℝ^3×H×W^, which is divided into a set of fixed-size image patches according to spots’ coordinates, denoted as *P* = {*p*_1_, …, *p*_*N*_}, where *N* is the number of spots.

### 2.3 Data preprocessing

Given an image patch *p*, we first sample a set of multi-view crops through uniform sampling, followed by random augmentations (e.g., color jitter, blur, stain jitter):

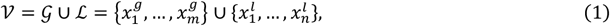

where 𝒢 and ℒ denote the sets of global and local crops, respectively. *m, n* are the numbers of crops. 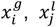 are subregions sampled from the patch *p*. Empirically, the area ratio of local crops ranges from 32% to 100%, whereas those of global crops ranges from 5% to 32%.

With the global crops, we further create masked global crops 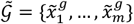 by sampling a random mask *ρ* ∈ {0,1}^*m*^ according to a masking ratio *r*.

### 2.4 Pretraining architecture

SpaFoundation is pretrained using a tailored iBOT^30^ framework to learn generalizable histological representations for inferring spatial gene expression (Fig. 1). Specifically, we construct a Vision Transformer (ViT)-based teacher-student network to learn patch-level representations which can be applied for spatial gene expression inference, as well as other downstream tasks, such as tumor detection and spatial clustering.

**Fig. 1.**
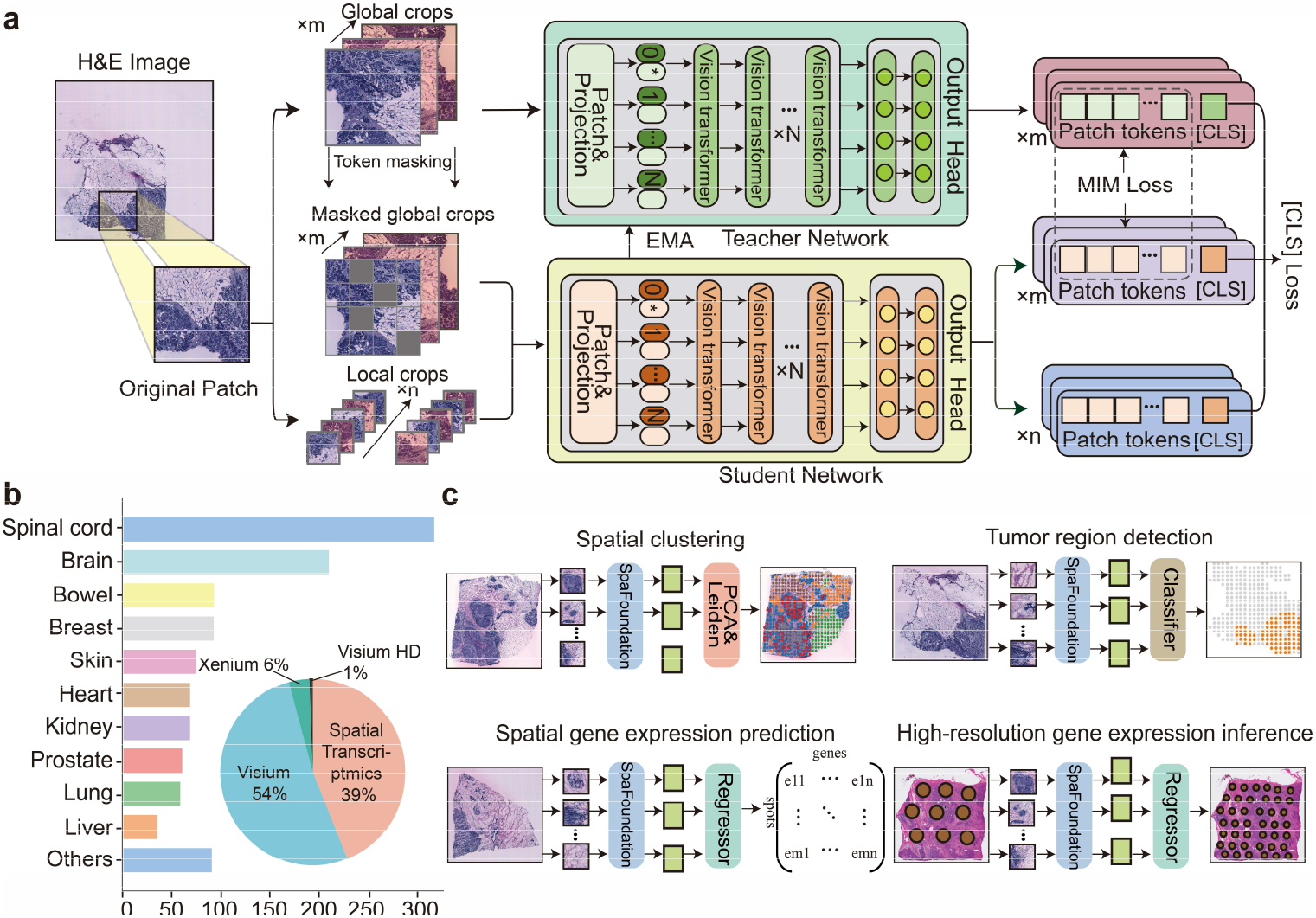
Pretraining of spafoundation and downstream task overview. (a) pretraining workflow of spafoundation, a teacher-student vision transformer architecture with self-distillation and masked image modeling (mim). (b) statistics of hest-1k-3 dataset, including 1,113 samples from 26 organ types and three technologies. (c) downstream tasks, including spatial clustering, tumor region detection, spatial gene expression prediction and high-resolution inference.

#### 2.4.1 Vision Transformer Encoding

During pretraining, each image view of the local crop *x*^*l*^, the global crop *x*^*g*^ and the masked global crops 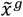 is independently encoded by a Vision Transformer (ViT) encoder. For a given crop, it is first divided into a sequence of fixed-size tokens. Each token is linearly projected into a d_1_-dimensional token embedding. A learnable class token *z*_*cls*_ is prepended to the token sequence, yielding the input representation 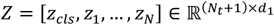, where *N*_*t*_ denotes the number of tokens.

The resulting token sequence is then processed by *L* transformer layers, each consisting of multi-head self-attention (MSA) and a feed-forward network (FFN) with residual connections and LayerNorm operations. Formally, for each layer *l* = 1, …, *L*, the transformation is formulated as:

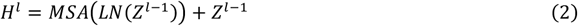

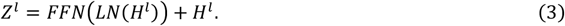

Within each attention head h, attention is defined as:

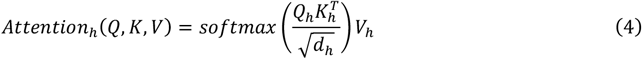

Where 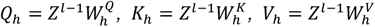, and *d*_*h*_ is the dimension per head. After training, the encoder outputs token representations, as well as class [CLS] representation *h*_*cls*_ which serves as the global representation of the crop.

#### 2.4.2 Teacher-student network

Both the teacher network and student network follow the same ViT encoding framework but maintain separate training parameters. The teacher network 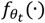 processes global crops and produces their class token representations 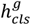. In parallel, the student network 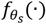 is applied to both local and masked global crops, yielding class [CLS] representations 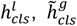, respectively. Formally, these class [CLS] representations are formulated as:

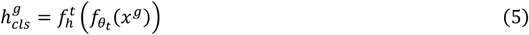

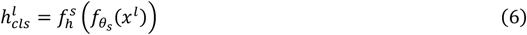

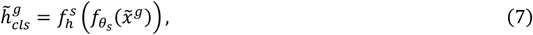

where 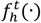 and 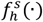 denote two-layer multi-layer perceptron (MLP) heads that project the token representations of different views into the same feature space.

### 2.5 Pretraining strategy

To efficiently capture both high-level global semantic information and fine-grained local structure features, we pretrain the model through two complementary objectives, i.e., self-distillation and masked image modeling (MIM). The self-distillation component enables the model to learn global semantic consistency across diverse image views, capturing high-level semantic representations. The MIM objective focuses on reconstructing masked regions, thereby encouraging the model to preserve detailed local morphological features. By jointly optimizing these two objectives, SpaFoundation facilitates comprehensive modeling of multi-level semantics in H&E images, resulting in biologically meaningful and robust feature representations. Next, we will elaborate on the components in detail.

#### 2.5.1 Self-distillation for learning high-level semantic features

Tissue structures often exhibit significant variability in shape, size, and orientation in the H&E images, posing significant challenges for learning generalizable representations. To circumvent the challenge and enhance semantic feature learning, we incorporate a self-distillation mechanism into the SpaFoundation model backbone. The self-distillation is a *discriminative* self-supervised objective that is designed to capture high-level semantic features by distilling knowledge from teacher to student. Specifically, as mentioned above, for an input image patch, we can obtain two disrupted views, i.e., the local crop *x^l^* and the global crop *x^g^*, via uniform sampling and random augmentations. These two views are then put through the teacher-student network to output [CLS] representations 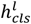 and 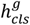, respectively. By minimizing the cross-entropy loss between the [CLS] representations of two views, SpaFoundation is encouraged to learn high-level semantic features. The self-distillation loss function ℒ_SD_ is defined as follows:

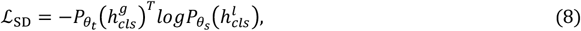

where P(.) denotes the probability distributions predicted by the teacher-student network. θ_t_ and θ_s_ refer to the parameters of the teacher and student networks, respectively.

#### 2.5.2 Masked image modeling for learning local semantic features

Building on BEiT^31^, we introduce masked image modeling (MIM) to enhance representation learning by predicting the visual tokens that are corresponding to the masked image patches. This component enables the model to capture local semantic features. Specifically, the teacher network takes as input the original global crops and outputs original patch tokens. The student network takes as inputs the masked global crops and outputs predicted patch tokens. The pre-training objective of MIM is to minimize the cross-entropy loss between the original and predicted patch tokens, guiding the student network to recover masked image information and thereby enhancing its ability to capture local semantic features. Formally, the MIM loss function ℒ_MIM_ is defined as follows:

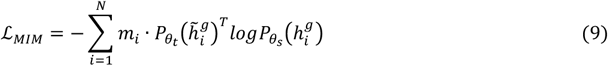

where N is the number of patches and m ∈ [0,1] denotes binary mask vector, with m_i_ = 1 indicating that the *i*-th patch of the image is masked, 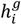 and 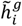 denote the i-th patch token from the teacher and student networks, respectively. P(·) denotes the probability distribution.

#### 2.5.3 Model optimization

The pretraining model is optimized jointly by the self-distillation loss ℒ_*SD*_ and the MIM loss ℒ_MIM_. The overall loss is defined as follows:

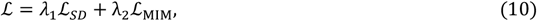

where *λ*_1_ and *λ*_2_ are weighting factors measuring the contributions of the self-distillation and MIM losses. During pretraining, the parameters of the student network *θ*_*s*_ are Exponentially Moving Averaged (EMA) to the parameters of the teacher network *θ*_*t*_.

## 3 Results

### 3.1 SpaFoundation enables accurate spatial domain segmentation using only image features

To assess the biological relevance of SpaFoundation’s learned histological features, we first evaluated its applicability to spatial domain segmentation using eight manually annotated human breast cancer tissues from the HER2+ dataset^28^. We compared SpaFoundation with three state-of-the-art non-spatial and spatial clustering methods, including Scanpy^32^, SpaGCN^33^, and stLearn^34^, under zero-shot scenario in which spot-level histological representations were extracted directly from the pretrained SpaFoundation model without fine-tuning. The resulting representations were subsequently subjected to PCA for dimensionality reduction and Leiden clustering.

We performed SpaFoundation and baseline methods on the H1 sample, which are manually annotated as six tissue regions, including invasive cancer, breast glands, in situ cancer, adipose tissue, immune infiltrate, and connective tissue (Fig. 2a). Both SpaFoundation and stLearn accurately identified invasive and in situ cancer regions with clear boundaries, whereas Scanpy and SpaGCN struggled with distinguishing these cancer types (Fig. 2b). SpaFoundation was the only method to accurately delineate adipose tissue with minimal noise. For the C1 sample which was annotated into three tissue regions, namely invasive cancer, adipose tissue, and connective tissue (Fig. 2c), we constrained all methods to produce the same number of clusters as the ground truth. SpaFoundation yielded spatial segmentations that align most closely with expert-defined histological annotations, while Scanpy, SpaGCN, and stLearn produced mixed or noisy boundaries (Fig. 2d). Notably, SpaFoundation clearly separated invasive cancer from connective tissue regions. Furthermore, we quantitatively evaluated the performance of different methods using 6 commonly used clustering metrics, including ARI, NMI, V-measure, Homo, and Mutual_Info. SpaFoundation consistently achieved the best performance across both samples and all measurement metrics (Fig. 2e, f). The boxplot across 8 samples showed that SpaFoundation achieved the highest median scores 0.3177, 0.3762, 0.3656, 0.3762, 0.3984, and 0.3566 of ARI, NMI, AMI, V-measure, Homogeneity, and Mutual Information, outperforming the second-best method stLearn by 30.00%, 30.81%, 32.80%, 30.81%, 22.42% and 28.96%, respectively (Fig. 2g). These results demonstrated SpaFoundation’s strong zero-shot performance, underscoring its ability to capture biologically relevant features from H&E images.

**Fig. 2.**
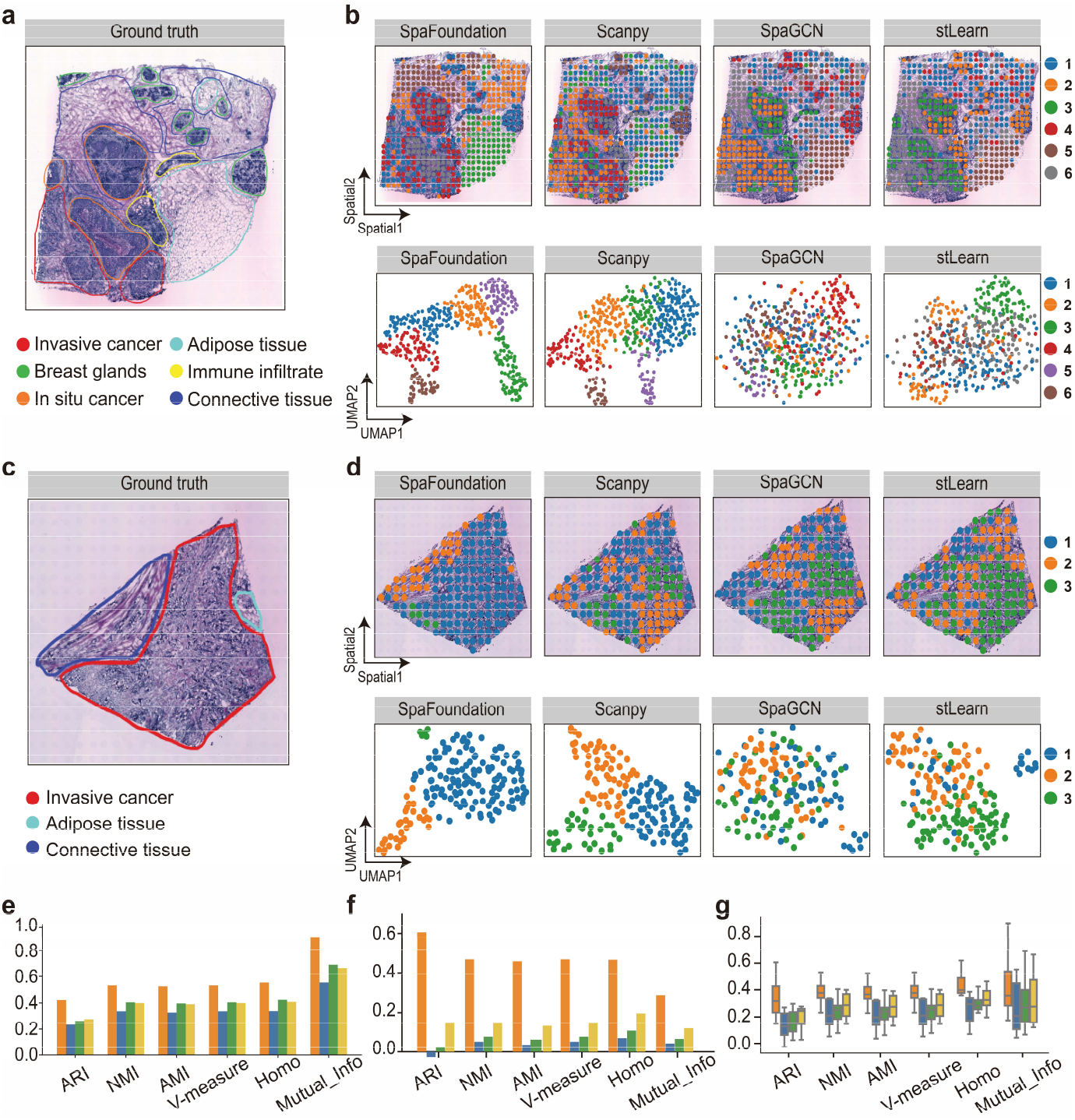
Performance comparison of SpaFoundation and scanpy, SpaGCN, and stLearn on the HER2+ dataset. (a) Ground truth of the H1 sample. (b) Spatial and UMAP distributions of SpaFoundation and baseline methods on the H1 sample. (c) Ground truth of the C1 sample. (d) Spatial and UMAP distributions of SpaFoundation and baseline methods on the C1 sample. (e)-(f) Quantitative evaluations of different methods in terms of ARI, NMI, V-measure, Homo, and Mutual_Info on the H1 and C1 samples, respectively. (g) Boxplots of different methods across 8 HER2+ samples evaluated using 6 metrics.

### 3.2 SpaFoundation accurately identifies tumor regions using pretrained histological representations

To further examine SpaFoundation’s ability to extract biologically meaningful features from tissue morphology, we applied it to tumor detection using the HBCIS dataset, a human breast cancer dataset comprising 68 tissue sections from 23 patients profiled by Spatial Transcriptomics^15^. We compared SpaFoundation with ResNet-in1k^24^ and VGG-in1k^35^, two widely used deep learning models pretrained on ImageNet-1K, and evaluated their performance using cross-validation. SpaFoundation was fine-tuned on data from 18 randomly selected patients and evaluated on the reminding 5 patients.

It can be observed that the tumor segmentation maps generated by SpaFoundation align more closely with the ground truth annotations and contained substantially less noise, whereas ResNet-in1k and VGG-in1k frequently misclassified non-tumor regions as tumor tissue (Fig. 3a). Quantitative evaluation further confirmed the advantage of SpaFoundation, which achieved the highest scores across Accuracy, Precision, F1 Score, and AUC Score (Fig. 3b). Notably, SpaFoundation reached an AUC of 0.94 across all test samples, surpassing ResNet-in1k by 4.16% and VGG-in1k by 1.55%. These results again demonstrate SpaFoundation’s strong ability to preserve biologically meaningful features from histological images. Importantly, the superior performance over conventional vision networks underscores the benefit of pretraining on domain-specific data, which enables the model to capture more informative and task-relevant histological representations.

**Fig. 3.**
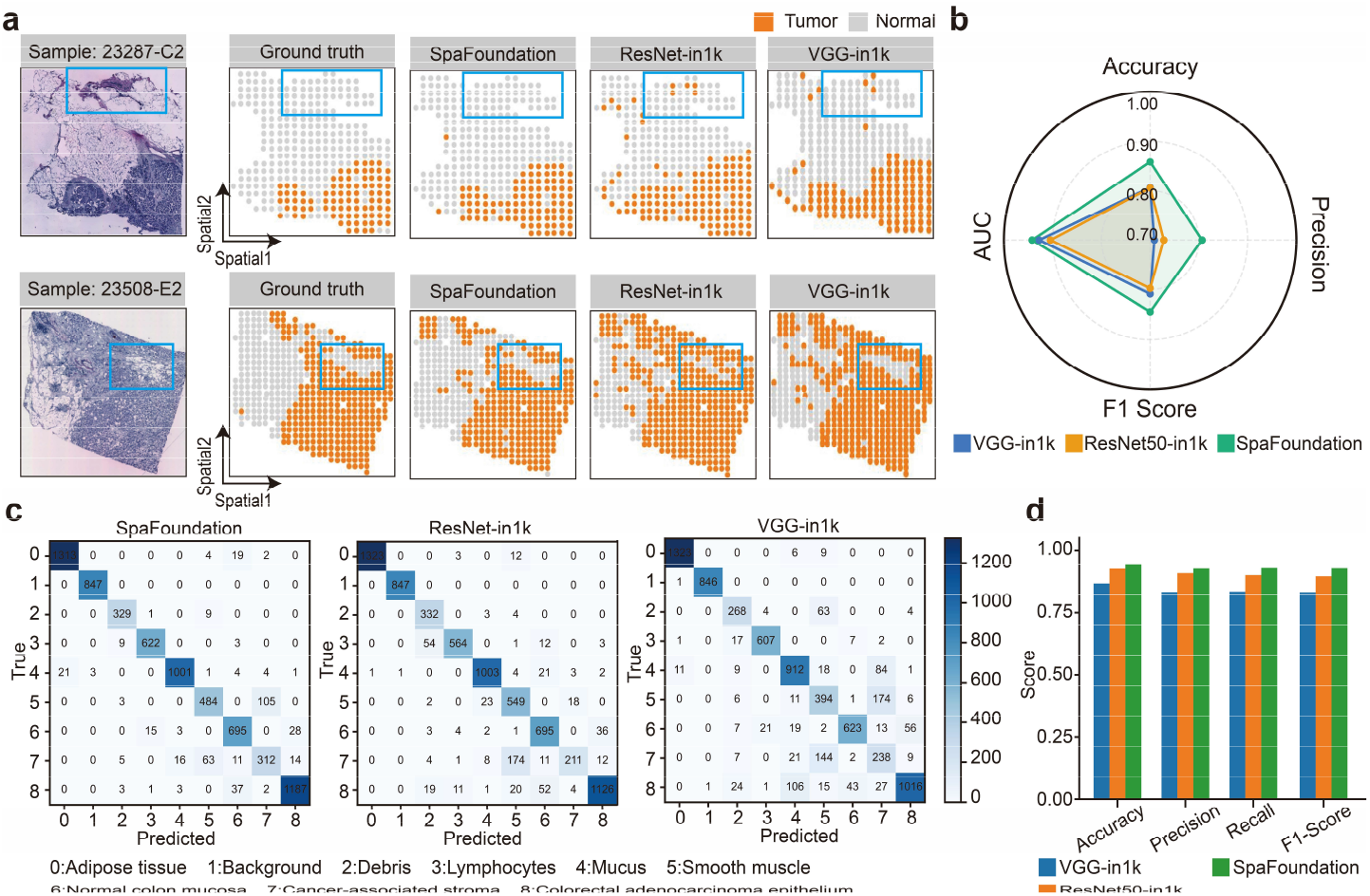
Performance comparison of SpaFoundation and ResNet-in1k and VGG-in1k on 23287-C2 and 23508-E2 samples from the HBCIS dataset for tumor detection. (a) H&E images and spatial maps of tumor and non-tumor regions identified by different methods on these two samples. (b) Quantitative evaluation on the HBCIS dataset in terms of metrics AUC, Accuracy, Precision, and F1-score. (c) Confusion matrices of SpaFoundation, ResNet-in1k, and VGG-in1k on the CRC-VAL-HE-7K dataset.(d) Quantitative evaluation on the CRC-VAL-HE-7K dataset in terms of metrics Accuracy, Precision, Recall, and F1-score.

Besides spatial transcriptomics H&E images, our SpaFoundation also demonstrates strong transferability to pathological imaging data. To evaluate this, we employed two colorectal cancer pathology datasets NCT-CRC-HE-100K, a human colorectal cancer tissue sample, and CRC-VAL-HE-7K, a human colorectal adenocarcinoma tissue sample, curated by Kather et al. These datasets contain 100,000 and 7,180 non-overlapping image patches, respectively, and share 9 tissue categories, including adipose tissue, background, debris, lymphocytes, mucus, smooth muscle, normal colon mucosa, cancer-associated stroma, and colorectal adenocarcinoma epithelium. We fine-tuned the pretrained SpaFoundation model on NCT-CRC-HE-100K and evaluated its performance on CRC-VAL-HE-7K. The confusion matrices indicated that SpaFoundation yields the most concentrated diagonal signals and minimal off-diagonal errors across 9 tissue types, outperforming ResNet-in1k and VGG-in1k, particularly in cancer-related types such as cancer-associated stroma and colorectal adenocarcinoma epithelium (Fig. 3c). Quantitative evaluation further validated the observations (Fig. 3d). These results further confirmed SpaFoundation’s ability to capture biologically meaningful features from histological images and demonstrated its strong transferability to pathological images.

### 3.3 SpaFoundation accurately predicts spatial gene expression profiles with histology images alone

Next, we evaluated SpaFoundation’s capability to infer spatial gene expression from histology images. We compared it with five state-of-the-art methods that are task-specific lightweight deep learning models, including HistoGene^18^, His2ST^17^, THItoGene^19^, BLEEP^20^, and TRIPLEX^21^. We used two independent cSCC^29^ and HER2+^28^ datasets for evaluation. The cSCC^29^ dataset consists of 12 squamous cell carcinoma samples from 4 patients, while the HER2+^28^ dataset contains 36 breast cancer samples from 8 patients. We comprehensively assess the model’s prediction capability using histological images under both highly expressed genes (HEG) and highly variable genes (HVG) settings.

Visually, the spatial expressions of selected genes, COL6A2, SPINK5, CD24, HMGB2, predicted by SpaFoundation more closely match the ground truth compared to baseline methods (Fig. 4), demonstrating the advantages of foundation model pretrained on large-scale H&E images over traditional lightweight models trained on specific individual sample. For better comparison, we quantitatively evaluated the performance of these methods using metrics PCC and SCC. Across both datasets and evaluation settings, SpaFoundation consistently achieved the highest improvements in both PCC and SCC, outperforming all baseline methods by a large margin (Table 1). Importantly, our performance advantage is particularly pronounced under more challenging HVG setting, where robust genes’ morphology feature extraction is critical. Moreover, it is observed that SpaFoundation exhibited stable and robust performance in all cases, while baseline methods exhibited sensitivities against the setting of HEG and HVG. For example, His2ST performed better on HVG prediction than on HEG, while BLEEP performed better on HEG than on HVG. These results validated SpaFoundation’s strong capability to accurately predict spatial gene expression directly from histological images.

**Table 1.** Comparison performance of different methods under HEG and HVG settings on cSCC and HER2+ datasets. Bold indicates the best performance, and underlined values indicate the second-best performance.

| Model | cSCC_HEG |  | cSCC_HVG |  | HER2+_HEG |  | HER2+_HVG |  |
| --- | --- | --- | --- | --- | --- | --- | --- | --- |
|  | PCC | SCC | PCC | SCC | PCC | SCC | PCC | SCC |
| HistoGene | 2.49% | 1.77% | 5.05% | 5.30% | 9.28% | 6.31% | 4.13% | 4.20% |
| His2ST | 11.67% | 10.92% | 11.72% | <u>13.34%</u> | 3.19% | 2.79% | 6.64% | <u>7.03%</u> |
| THItoGene | 3.31% | 1.90% | -1.15% | 0.08% | 1.12% | 0.65% | 0.97% | 1.12% |
| BLEEP | <u>16.22%</u> | <u>15.76%</u> | <u>12.06%</u> | 12.62% | <u>11.51%</u> | <u>9.44%</u> | 6.56% | 6.30% |
| TRIPLEX | 13.84% | 12.63% | 6.05% | 6.48% | 10.16% | 7.98% | <u>7.17%</u> | 7.50% |
| SpaFoundation-R | 14.76% | 12.22% | 10.12% | 11.70% | 14.00% | 9.33% | 8.62% | 8.83% |
| SpaFoundation | **18.82%** | **18.28%** | **13.08%** | **14.49%** | **15.56%** | **12.42%** | **9.86%** | **10.09%** |

**Fig. 4.**
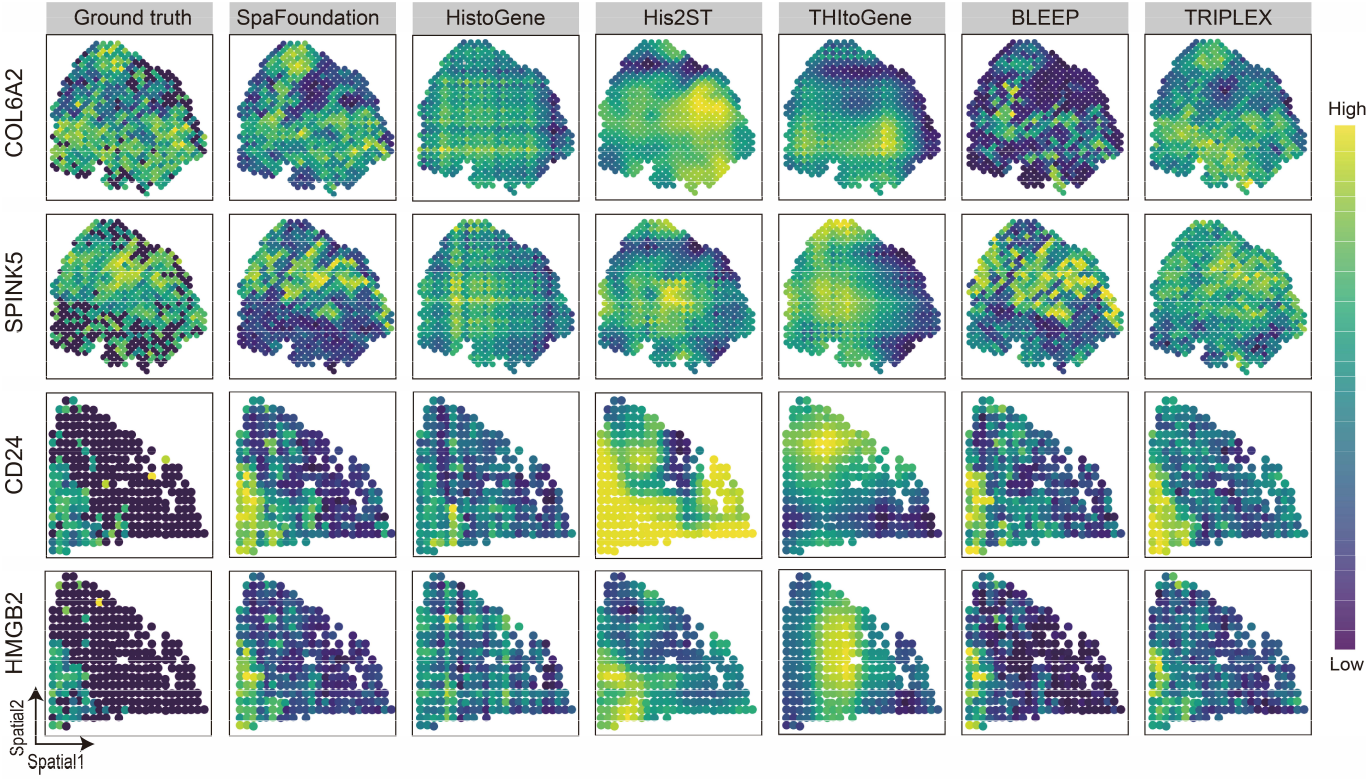
Spatial maps of predicted gene expression across the cSCC and HER2+ datasets generated by SpaFoundation, HistoGene, His2ST, THItoGene, BLEEP, and TRIPLEX.

The comparison between SpaFoundation with its variant SpaFoundation-R, in which the model parameters were randomly initialized instead of using pretraining parameters, indicated that SpaFoundation consistently achieved higher scores than SpaFoundation-R across both settings and both evaluation metrics (Table 1). These results highlight the critical contributions of pretraining to accurate spatial gene expression inference.

### 3.4 SpaFoundation enables accurate inference of high-resolution gene expression from low-resolution histology images

We further evaluated SpaFoundation’s ability to infer high-resolution spatial gene expression from low-resolution spatial transcriptomics (ST) data. ST technologies often face a trade-off between spatial resolution and transcriptome coverage, with imaging-based platforms offering high resolution but limited gene coverage, while sequencing-based platforms capture more genes at lower resolution. To address this, we applied SpaFoundation on sequence-based ST data to infer high-resolution gene expression.

Specifically, we used a human breast cancer Visium HD dataset with 16μm and 8μm spatial resolutions. We fine-tuned the pre-trained SpaFoundation model on 16μm patches (∼140,000 spots) and tested it on 8μm patches (∼530,000 spots). We compared SpaFoundation to ST-Net^15^, a state-of-the-art method for high-resolution gene prediction. The visualization results showed that SpaFoundation more faithfully recapitulated ground-truth spatial gene-expression patterns for the selected genes ERBB2 and DMBT1 (Fig. 5a). Quantitatively, we evaluated the prediction accuracy of top 1,000 highly variable genes (HVGs) using measure metrics PCC, SCC, MSE, and RMSE. As a result, SpaFoundation achieved median PCC, SCC, MSE, and RMSE of 0.185, 0.1683, 0.1152, and 0.3394, improving over ST-Net by 151.02%, 9.5%, 48.41%, and 28.18%, respectively (Fig. 5b). These results highlight SpaFoundation’s superior capability in inferring high-resolution gene expression using low-resolution ST data.

**Fig. 5.**
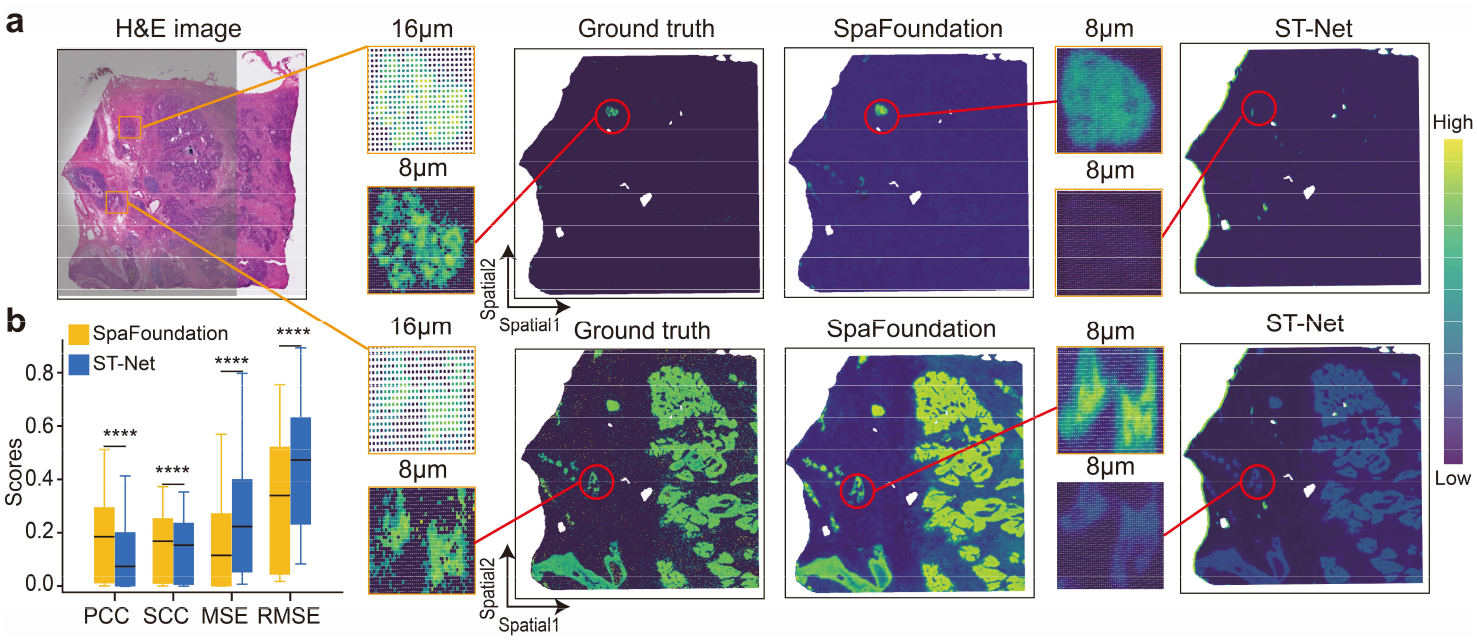
Comparison of SpaFoundation with ST-Net in high-resolution gene expression inference: (a) H&E image and spatial maps of genes ERBB2 (top) and DMBT1 (bottom) predicted by SpaFoundation and ST-Net. (b) Quantitative comparison of SpaFoundation and ST-Net in terms of metrics PCC, SCC, MSE, and RMSE.

## 4 Discussion

In this study, we present SpaFoundation, a histology-driven foundation model tailored to advance spatial gene expression prediction. Unlike traditional task-specific lightweight models, SpaFoundation can accurately infer spatial gene expression without requiring additional retraining when encountering new sample. To the best of our knowledge, SpaFoundation is among the first foundation models explicitly pretrained on large-scale H&E images for spatial gene expression inference. Although recent efforts have explored using existing pretrained vision foundation models to extract image features for gene expression prediction, their performance remains limited. A major limitation lies in the mismatch between their pretraining domains, typically pathological images, and the H&E images derived from spatial transcriptomics. This domain gap restricts their ability to capture the morphological features most relevant to spatial gene expression. To circumvent the challenge, we curated HEST-1K, a large-scale dataset comprising 1,113 samples spanning 26 tissue organs, each consisting of paired H&E image and spatial transcriptomics. Building upon the iBOT (image BERT pre-training with Online Tokenizer) framework, we pretrained SpaFoundation on this domain-specific dataset to learn general-purpose histological representations by modeling the intrinsic dependencies between image patches. The integration of self-distillation and masked image modeling (MIM) enables the model to simultaneously capture fine-grained local morphology and high-level semantic structure, making it possible to capture more biologically meaningful features. Evaluated across 117 samples from multiple ST platforms, SpaFoundation demonstrated strong zero-shot and few-shot performance across diverse downstream tasks, highlighting its versatility as a histology-driven foundation model for spatial transcriptomics.

The superior performance of SpaFoundation can be attributed primarily to two aspects. First, the combination of self-distillation and masked image modeling (MIM) fosters more expressive and hierarchical representation learning. Second, SpaFoundation is pretrained on spatial transcriptomics-specific H&E images, ensuring that the learned features are highly aligned with spatial gene expression. Despite these advantages, our method still faces limitations. The HEST-1K dataset, though substantial, remains modest relative to the scale typically required for fully exploiting the capacity of foundation models. As more large-scale, high-quality H&E datasets become available, we expect SpaFoundation to further benefit from broader and more diverse pretraining, ultimately leading to more robust and accurate spatial gene expression inference.

## Notes

### Competing Interest Statement

The authors have declared no competing interest.

### Summary of Updates

Some experimental results and analytical content have been updated.

### Funding

- National Natural Science Foundation of China under Grant: No.62225209, Hunan Provincial Graduate Student Scientific Research Innovation Project, CX20250238

## References

1. Yayon, N. et al. A spatial human thymus cell atlas mapped to a continuous tissue axis. Nature 635, 708–718 (2024).

2. Vannan, A. et al. Spatial transcriptomics identifies molecular niche dysregulation associated with distal lung remodeling in pulmonary fibrosis. Nat. Genet. 57, 647–658 (2025).

3. Polonsky, M. et al. Spatial transcriptomics defines injury specific microenvironments and cellular interactions in kidney regeneration and disease. Nat. Commun. 15, 7010 (2024).

4. Li, L. et al. An organ-wide spatiotemporal transcriptomic and cellular atlas of the regenerating zebrafish heart. Nat. Commun. 16, 3716 (2025).

5. Qian, X. et al. Spatial transcriptomics reveals human cortical layer and area specification. Nature 1–11 (2025) doi:10.1038/s41586-025-09010-1.

6. Yang, W. et al. Deciphering cell–cell communication at single-cell resolution for spatial transcriptomics with subgraph-based graph attention network. Nat. Commun. 15, 7101 (2024).

7. Zhang, J. et al. Spatiotemporally resolved transcriptomics reveals the cellular dynamics of human retinal development. Nat. Commun. 16, 2307 (2025).

8. Feng, J. et al. High-resolution spatial transcriptomics and cell lineage analysis reveal spatiotemporal cell fate determination during craniofacial development. Nat. Commun. 16, 4396 (2025).

9. Wang, X. et al. Three-dimensional intact-tissue sequencing of single-cell transcriptional states. Science (2018) doi:10.1126/science.aat5691.

10. Stickels, R. R. et al. Highly sensitive spatial transcriptomics at near-cellular resolution with Slide-seqV2. Nat. Biotechnol. 39, 313–319 (2021).

11. Chen, A. et al. Spatiotemporal transcriptomic atlas of mouse organogenesis using DNA nanoball-patterned arrays. Cell 185, 1777-1792.e21 (2022).

12. Takei, Y. et al. Integrated spatial genomics reveals global architecture of single nuclei. Nature 590, 344–350 (2021).

13. Liu, J. et al. Concordance of MERFISH spatial transcriptomics with bulk and single-cell RNA sequencing. Life Sci. Alliance 6, (2023).

14. He, S. et al. High-plex imaging of RNA and proteins at subcellular resolution in fixed tissue by spatial molecular imaging. Nat. Biotechnol. 40, 1794–1806 (2022).

15. He, B. et al. Integrating spatial gene expression and breast tumour morphology via deep learning. Nat. Biomed. Eng. 4, 827–834 (2020).

16. Li, S., Gai, K., Dong, K., Zhang, Y. & Zhang, S. High-density generation of spatial transcriptomics with STAGE. Nucleic Acids Res. 52, 4843–4856 (2024).

17. Zeng, Y. et al. Spatial transcriptomics prediction from histology jointly through Transformer and graph neural networks. Brief. Bioinform. 23, bbac297 (2022).

18. Pang, M., Su, K. & Li, M. Leveraging information in spatial transcriptomics to predict super-resolution gene expression from histology images in tumors. Preprint at 10.1101/2021.11.28.470212 (2021).

19. Jia, Y., Liu, J., Chen, L., Zhao, T. & Wang, Y. THItoGene: a deep learning method for predicting spatial transcriptomics from histological images. Brief. Bioinform. 25, bbad464 (2023).

20. Xie, R. et al. Spatially Resolved Gene Expression Prediction from H&E Histology Images via Bi-modal Contrastive Learning. NeurIPS (2023).

21. Chung, Y., Ha, J. H., Im, K. C. & Lee, J. S. Accurate Spatial Gene Expression Prediction by integrating Multi-resolution features. Preprint at http://arxiv.org/abs/2403.07592 (2024).

22. Fu, X. et al. Spatial gene expression at single-cell resolution from histology using deep learning with GHIST. Nat. Methods 22, 1900–1910 (2025).

23. Wang, Y. et al. FmH2ST: foundation model-based spatial transcriptomics generation from histological images. Nucleic Acids Res. 53, gkaf865 (2025).

24. He, K., Zhang, X., Ren, S. & Sun, J. Deep Residual Learning for Image Recognition. in 2016 IEEE Conference on Computer Vision and Pattern Recognition (CVPR) 770–778 (IEEE, Las Vegas, NV, USA, 2016). doi:10.1109/CVPR.2016.90.

25. Chen, R. J. et al. Towards a general-purpose foundation model for computational pathology. Nat. Med. 30, 850–862 (2024).

26. Xue, S., Wang, C., Fan, X. & Min, W. Inferring Super-Resolved Gene Expression by Integrating Histology Images and Spatial Transcriptomics with HISTEX.

27. Jaume, G. et al. HEST-1k: A Dataset for Spatial Transcriptomics and Histology Image Analysis. Preprint at http://arxiv.org/abs/2406.16192 (2024).

28. Andersson, A. et al. Spatial deconvolution of HER2-positive breast cancer delineates tumor-associated cell type interactions. Nat. Commun. 12, 6012 (2021).

29. Ji, A. L. et al. Multimodal Analysis of Composition and Spatial Architecture in Human Squamous Cell Carcinoma. Cell 182, 497-514.e22 (2020).

30. Zhou, J. et al. iBOT: Image BERT Pre-Training with Online Tokenizer. Preprint at http://arxiv.org/abs/2111.07832 (2022).

31. Bao, H., Dong, L., Piao, S. & Wei, F. BEiT: BERT Pre-Training of Image Transformers. Preprint at 10.48550/arXiv.2106.08254 (2022).

32. Wolf, F. A., Angerer, P. & Theis, F. J. SCANPY: large-scale single-cell gene expression data analysis. Genome Biol. 19, 15 (2018).

33. Hu, J. et al. SpaGCN: Integrating gene expression, spatial location and histology to identify spatial domains and spatially variable genes by graph convolutional network. Nat. Methods 18, 1342–1351 (2021).

34. Pham, D. T., Tan, X., Xu, J., Grice, L. F. & Nguyen, Q. H. stLearn: integrating spatial location, tissue morphology and gene expression to find cell types, cell-cell interactions and spatial trajectories within undissociated tissues. bioRxiv (2020) doi:10.1101/2020.05.31.125658.

35. Simonyan, K. & Zisserman, A. Very Deep Convolutional Networks for Large-Scale Image Recognition. Preprint at 10.48550/arXiv.1409.1556 (2015).

